# The TERB1-TERB2-MAJIN complex of mouse meiotic telomeres dates back to the common ancestor of metazoans

**DOI:** 10.1101/755512

**Authors:** Irene da Cruz, Céline Brochier-Armanet, Ricardo Benavente

## Abstract

**Background:** Meiosis is essential for sexual reproduction, and generates genetically diverse haploid gametes from a diploid germ cell. Reduction of ploidy depends on active chromosome movements during early meiotic prophase I. Chromosome movements require telomere attachment to the nuclear envelope. This attachment is mediated by telomere adaptor proteins. Telomere adaptor proteins have to date been identified in fission yeast and mice. In the mouse, they form a complex composed of the meiotic proteins TERB1, TERB2, and MAJIN. No sequence similarity was observed between these three mouse proteins and the adaptor proteins of fission yeast, raising the question of the evolutionary history and significance of this specific protein complex.

**Result:** Here, we show the TERB1, TERB2, and MAJIN proteins are found throughout the Metazoa and even in early branching non-bilateral species such as Cnidaria, Placozoa and Porifera. Metazoan TERB1, TERB2, and MAJIN showed comparable domain architecture across all clades. Furthermore, the protein domains involved in the formation of the complex as well as those involved for the interaction with the telomere shelterin protein and the LINC complexes revealed high sequence similarity. Finally, gene expression in the cnidarian *Hydra vulgaris* provide evidence that the TERB1-TERB2-MAJIN complex is selectively expressed in the germ line.

**Conclusion:** Our results indicate that the TERB1-TERB2-MAJIN complex has an ancient origin in metazoans, suggesting conservation of meiotic functions.

## Background

Meiosis is a cell division mode during which one round of DNA replication is followed by two successive rounds of chromosome segregation leading, to the generation of haploid cells. During the prophase I stage, a series of specialized events including homologous chromosome pairing, synapsis, and recombination are crucial for the faithful completion of meiosis. These processes mainly depend on evolutionarily conserved chromosome movements and require the association of telomeres with the nuclear envelope (NE) (Hiraoka & Dernburg., 2009; Koszul & Kleckner, 2009).

The forces required for meiotic chromosome movements are generated in the cytoplasm and transduced into the nucleus via LINC (Linker of Nucleoskeleton and Cytoskeleton) complexes of the nuclear envelope (NE) (Alsheimer, 2009; Bhalla & Dernburg, 2008; Sheehan & Pwlowski, 2009; Woglar & Jantsch, 2014). In mice, this complex is composed of the transmembrane proteins SUN1 and SUN2 of the inner nuclear membrane (INM), and the KASH5 protein in the outer nuclear membrane (ONM) (Ding et al., 2007; Link et al., 2014; Horn et al., 2013; Morimoto et al., 2012; Schmitt et al., 2007). In addition, the meiotic telomere adaptor proteins TERB1, TERB2, and MAJIN (Daniel et al., 2014; Shibuya et al., 2014; 2015) provide the physical linkage of telomeres to the NE. To accomplish this, TERB1 interacts with SUN1 through its N-terminal ARM repeat domain and with the telomeric shelterin protein TRF1 through a region flanking its TERB2-binding site (Supplementary Fig. 1) (Long et al., 2017; Pendlebury et al., 2017; Shibuya et al., 2014; 2015; Zhang et al., 2017). Simultaneously, a specific C-terminal region of TERB2 interacts with the MAJIN N-terminal domain (Shibuya et al., 2015; Wang et al., 2019). MAJIN possesses DNA binding properties and is anchored to the INM via a transmembrane helix at its C-terminus (Shibuya et al., 2014; 2015; Dunce et al., 2018; Wang et al., 2019).

The SUN-KASH proteins, are widely conserved in animals, nematodes, yeast, and plants (Link & Jantsch, 2019; Pradillo et al., 2019; Zeng et al., 2018). However, the conservation of the meiotic specific telomere adaptor proteins is less clear. In yeast, a similar mechanism for anchoring telomeres to the NE has been identified. It involves the Bqt1 −2 meiotic telomere adaptor that connects, the Taz1-Rap1 telomere protein to Sad1 SUN domain protein in a complex with Btq3-4, INM proteins (Chikashige & Hiraoka, 2001; Chikashige et al, 2006; 2009; Miki et al., 2004; Cooper et al., 1998) (Supplementary Fig. S2). However, no protein sequence similarity these yeast telomere adaptor proteins and those of mice has been detected.

The high divergence between yeast and mice telomere adaptor proteins makes it unclear how conserved the mechanisms mediating the dynamic anchoring of meiotic telomeres to the NE. To probe the origin of the murine TERB1-TERB2-MAJIN complex, we used computational methods to identify their putative orthologues in public databases and subjected candidates to expression studies. Our results indicate that the TERB1-TERB2-MAJIN complex is evolutionarily ancient and that dates back to the common ancestor of metazoans.

## Results

### TERB1, TERB2 and MAJIN are ancestral metazoans proteins

To identify candidate TERB1, TERB2 and MAJIN homologues in other taxa, we carried out a bioinformatic screen of public sequence database using PSI-BLAST (Altschul et al., 1997). Our survey of public sequence databases identified homologues of the mouse meiotic telomere complex proteins TERB1, TERB2, and MAJIN mainly in metazoans (Supplementary Information, Table S1, S2 and S3). Most sequences were obtained from Deuterostomes and especially in the Vertebrata, but candidates were also readily identified in Cephalochordata, Echinodermata, and Hemichordata. Beside the deuterostomes, homologues were also detected in the Lophotrochozoa principally Mollusca, Annelida, and Brachiopoda. Only a few putative homologous were found among Ecdysozoans in the Priapulida and Arthropoda clades. We also identified putative homologues of TERB1, TERB2, and MAJIN in non-bilaterians such as Cnidaria, Placozoa, and Porifera. The taxonomic distributions of TERB1, TERB2, and MAJIN were very similar, meaning that all three corresponding genes are present in metazoan species. Of note, TERB1, TERB2, and MAJIN are present in a single copy in vertebrates, despite the two rounds of whole genome duplication that occurred during the evolutionary history of this group (Dehal & Boore, 2005). This indicates that paralogues resulting from these events were not retained during evolution. Similarly, no paralogues were observed in other metazoan lineages.

These results implied that these three proteins are likely ancient in metazoans, and arose before the emergence of bilaterians.

TERB1 multiple sequence alignments revealed high similarity in protein domain organization with N-terminal ARM repeats (aa 16-384) and C-terminal MYB (aa 715-747) domain shared between Porifera, Cnidaria (Hydrozoans), Annelida, Mollusca, Brachiopoda, Echinodermata (Asterozoa) and Vertebrata (Fig. 1A, Supplementary Fig.S3). Surprisingly, candidate TERB1 sequences from Cnidaria (Anthozoa), Arthropoda, Priapulida, Cephalopoda, Hemichordate and Cephalochordate appear to lack a Myb domain (Supplementary Information, Table S2). Nonetheless, closer inspection of the sequence alignment indicated amino acid conservation flanking the essential site for TERB2 binding (T2B) (aa 593-622 of mice) in TERB1 (Shibuya 2015; Long et al, 2017; Pendlebury et al., 2017; Zhang et al., 2017) (Supplementary Fig. S4). The probable absence of a Myb domain suggests either annotation errors or the loss of this domain during evolution. In addition, TERB1 interacts directly with the shelterin complex protein TRF1 via its TRFB domain, (aa 523-699 of mice) which is adjacent to the TERB2 binding site (T2B) near its C-terminus (Shibuya et al., 2014; 2015; Zhang et al., 2017; Long et al., 2017; Pendlebury et al., 2017). Previous biochemical and structural data have analyzed the interaction between TERB1 and TRF1 using a short peptide sequence (IxLxP, aa 646-650 of human) that highly resembles the TRF1-binding motif [F/YxLxP], of the shelterin-associated protein TIN2 (Chen et al., 2008; Long et al 2017; Wang el at 2018;). However, this motif was conserved mostly in Vertebrata (Supplementary Fig. S3).

**Fig. 1.**
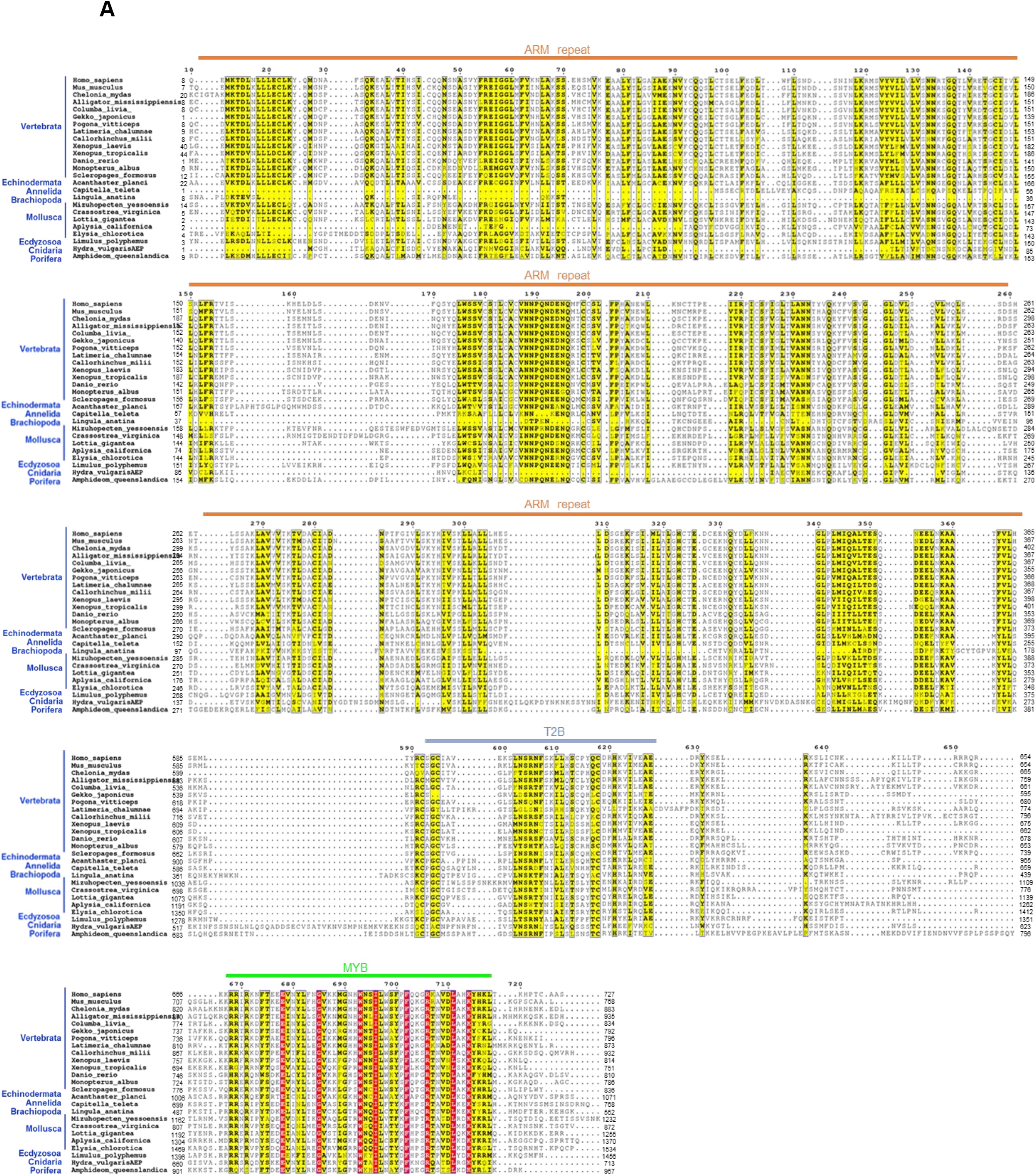
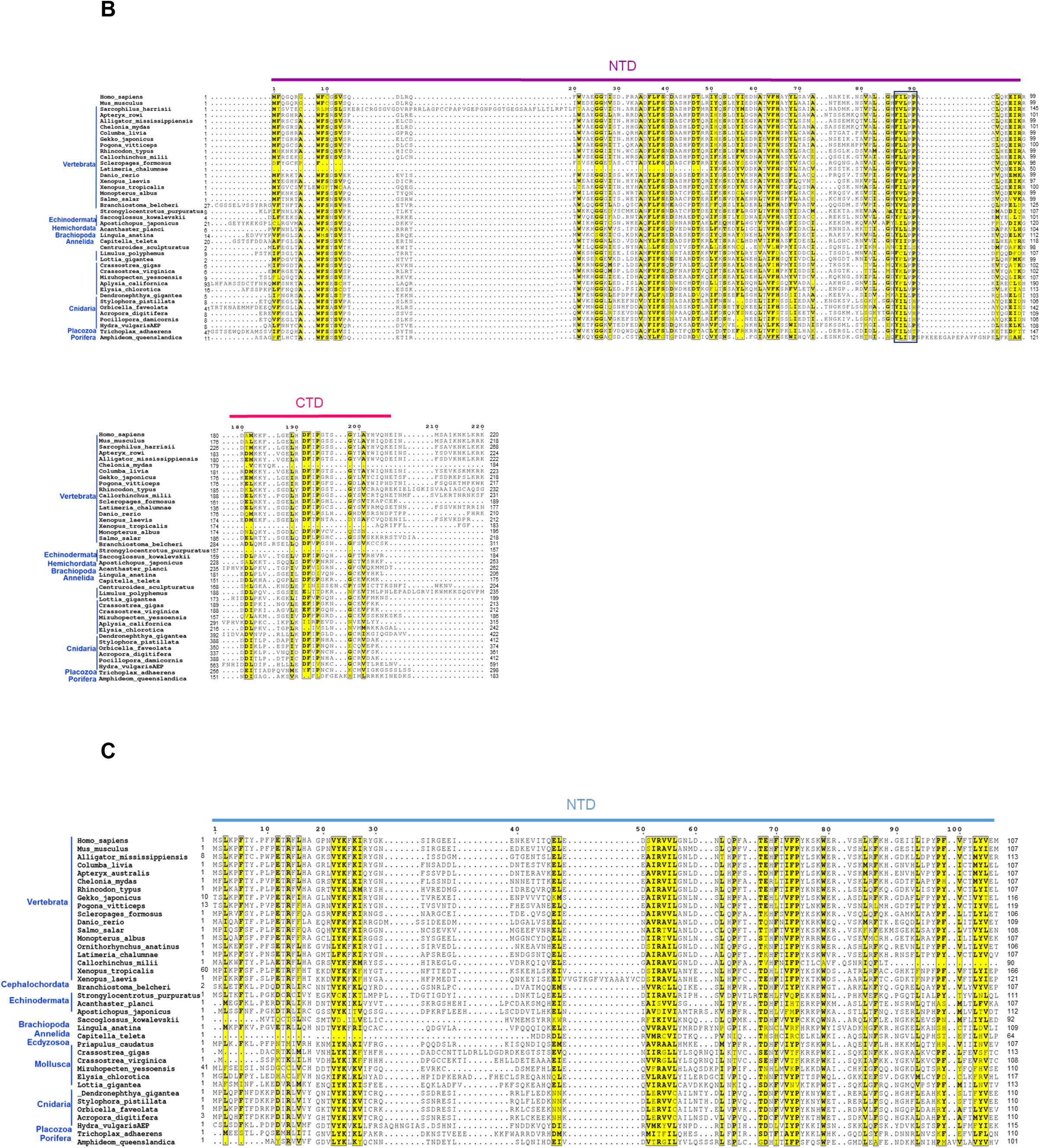
TERB1-TERB2-MAJIN proteins share domain architecture organization and functional sites across metazoans. Multiple sequence alignment of the most conserved regions of putative orthologs of mice TERB1 (A), TERB2 (B), and MAJIN (C) from various representative metazoan species with PROMALS3D and annotated using ESPrit 3.0. The threshold for grouping of the residues was set to 70% and is depicted in yellow. Amino acid positions conserved in all taxa are highlighted in red. Species were grouped according to their taxonomic group denoted with blue and the names of the species are shown at the left. The horizontal lines above the alignment indicate domain boundaries. Armadillo repeat; ARM repeat, T2B; binding site of TERB2 and MYB; homeodomain); N-terminal (NTD) and C-terminal (CTD) of TERB2 and N-terminal MAJIN (NTD). The motif [F/YxLxP] detected in all TERB2 sequences is shown in the dark blue rectangle.

The multiple alignment of TERB2 sequences revealed prominent conservation stretches spanning the entire N-terminal domain (aa 1-116 of mouse) (Fig. 1B). Furthermore, smaller stretches along the C-terminal region of TERB2 (aa 174-209 of mice) were found to be conserved among taxa (Fig. 1B, Supplementary Fig. S5). This result indicates that the most conserved features in TERB2 comes from the protein regions essential for the interaction with TERB1 and with MAJIN (Shibuya et al., 2015; Dunce et al., 2018; Wang et al., 2019). Unexpectedly, at the most end of the N-terminal domain (human and mouse aa 86-90) we noted in all the taxa a motif equivalent to [F/YxLxP], involved in the binding of shelterin-associated proteins to the TRFH domain of TRF1 and TRF2 (Fig. 1B) (Chen et at., 2008). This finding suggests that telomeric shelterin proteins TRF1 could potentially recruit TERB2 in addition to TERB1 (see above).

Finally, a multiple sequence alignment of MAJIN candidates showed sequence similarity restricted to the N-terminus (aa 2-194 of mice) (Fig. 1C, Supplementary Fig S6). This region interact with the C-terminal part of TERB2 (Shibuya et al., 2015; Wang et al., 2019). The transmembrane domain close to the C-terminus of murine MAJIN (aa 252-256) appears to be divergent among the taxa (Supplementary, Fig. S6).

Overall, the multiple sequence alignment of putative TERB1-TERB2-MAJIN orthologues showed comparable protein organization to the mouse proteins and revealed conserved stretches of high similarity in those domains required for the interaction with binding partners. These findings support the notion that despite their great divergence, these proteins are bona fide homologous, meaning that they derive from a common ancestor.

To recover the evolutionary history of TERB1, TERB2 and MAJIN proteins, taxonomically balanced Bayesian trees were built (Fig 2 A-C). The deepest nodes of the trees (especially TERB2 and MAJIN) are poorly resolved (weak posterior probabilities < 0.75), probably due to the relatively small number of sites that were kept for the phylogenetic analyses and indicating lack of phylogenetic signal rather than a true conflicting signal (Fig. 2 A-C). Nevertheless, it is noticeable that within Vertebrata, the relationship among sequences are consistent with the systematics, and that most Lophotrochozoa sequence group together (Fig. 2 A-C). In the Ecdysozoa clade, candidate proteins that belong to Arthropoda (*Zootermopsis nevadensis* and *Centruroides sculpturatus*) for TERB1 and TERB2 are less conserved and showed relatively long branches indicating suggesting a fast rate of evolution (Fig. 2A-B).

**Fig. 2.**
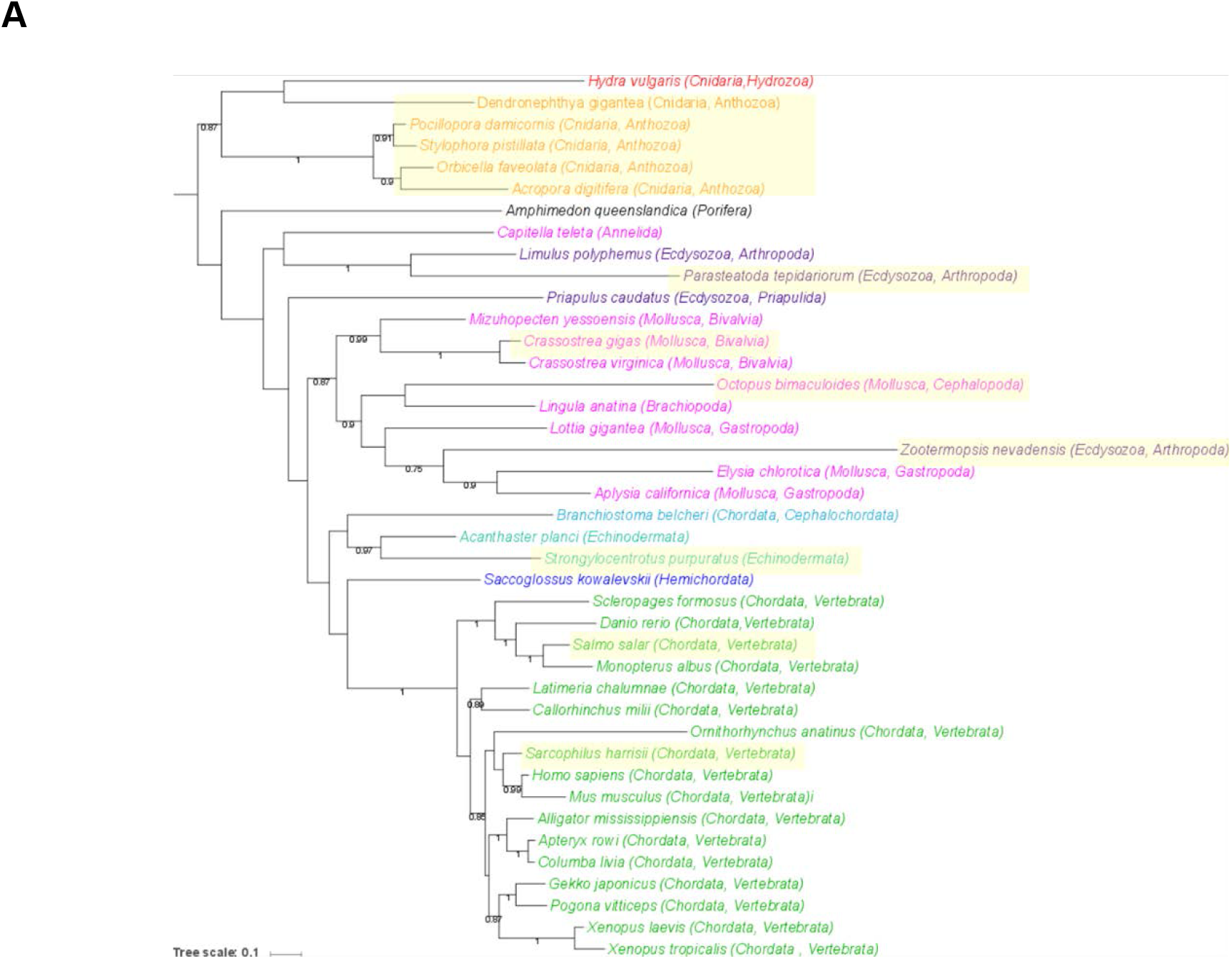
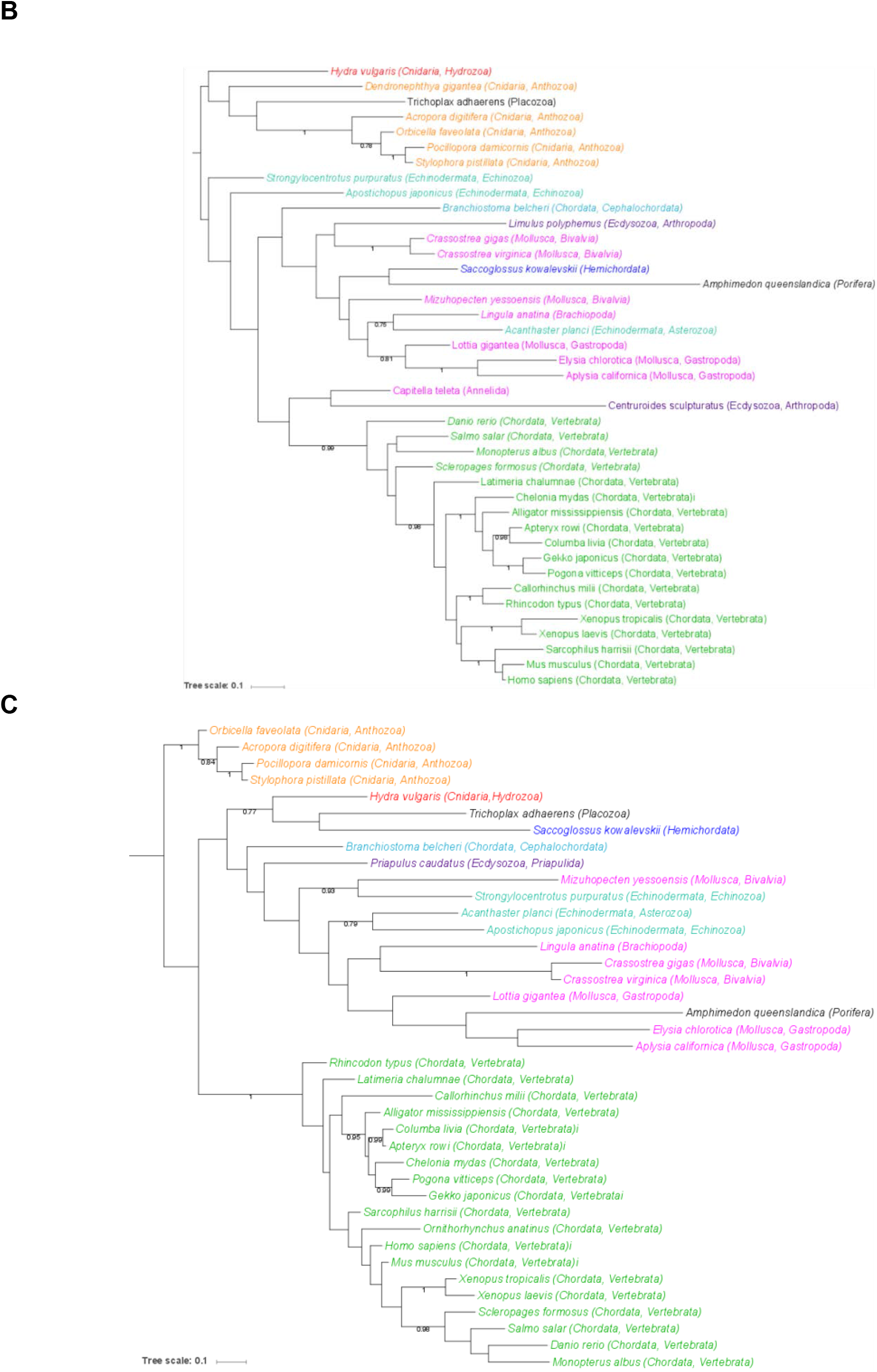
Evolutionary history of metazoan TERB1-TERB2-MAJIN proteins. Unrooted Bayesian tree of TERB1 (A), TERB2 (B) and MAJIN (C) proteins inferred with MrBayes. Numbers at branches represent posterior probabilities. For clarity, values of < 0.75 were omitted from the tree. The taxa lacking a MYB domain for TERB1 are highlighted in yellow. The length of each branch is proportional to the number of amino acid substitutions per site that have occurred. The bar displays the average number of substitutions per site (scale at top left of each panel).

In summary, these results clearly indicate that MAJIN, TERB1, and TERB2 were present in the ancestor of all present-day metazoans and are conserved in most present-day lineages including the early diverging Porifera, Placozoa, and Cnidaria (Fig. 2A-C).

### Expression of TERB1, TERB2 and MAJIN in the basal metazoan *Hydra*

According to our analysis, we have identified orthologues of TERB1, TERB2, and MAJIN across metazoans. To provide insight into a possible meiotic role of the candidate orthologues in species evolutionarily distant from the mouse, we decided to investigate the expression pattern of putative TERB1, TERB2 and MAJIN orthologues in the basal metazoan *Hydra vulgaris*. Equal amounts of total RNA were isolated from four different body regions: head, body column, testes, and foot TERB1, TERB2, and MAJIN transcript were detected by RT-PCR using specific primers that spanned exon-exon junctions in the predicted transcript of *Hydra vulgaris* (Supplementary Information, Table S4).

Our experiments showed that putative Hydra Terb1, Terb2 and Majin were specifically detected in the testis fraction. Faint signals in the body column fraction result from slight contamination with the previously associated testis tissue. Amplification of Hydra actin demonstrated the equal concentration of mRNA in the different fractions (Fig. 3A).

**Fig. 3.**
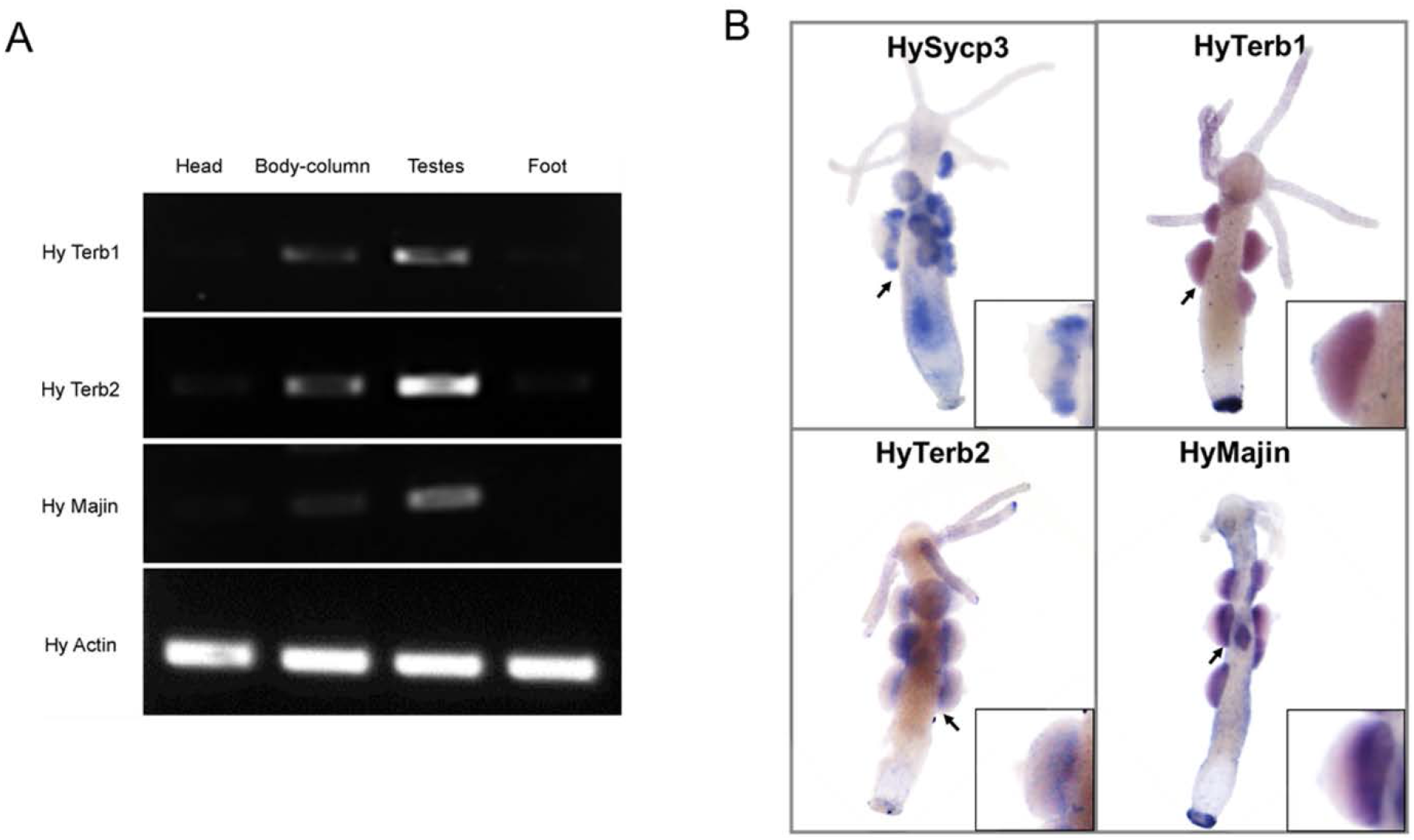
Gonad-specific expression Terb1, Terb2, Majin and Actin in *Hydra vulgaris*. (**A**) Terb1, Terb2, Majin were detected in Hydra tissues by RT-PCR. Actin was used as a control. All three transcripts are exclusively found in the testes. (**B**) Whole mount in situ hybridization to detect HySycp3, HyTerb1, HyTerb2, and HyMajin transcript. Arrows indicate the regions shown enlarged in the insets. All four transcripts were detected in cells at the base of the testes.

To confirm this expression pattern and to evaluate the spatial localization of Hydra Terb1, Terb2, and Majin transcripts we performed whole mount in situ hybridization (WMIH) (Fig. 3B). Sycp3, a specific marker of meiotic cells, was used as a positive control (Fraune et al. 2012). Transcript signals corresponding to Terb1, Terb2, Majin and Sycp3 were restricted to the basal layer of the testes, i.e. the site where meiotic cells are located (Kuznetsov et al., 2001; Fraune et al., 2013).

In addition, we achieved to amplify the complete coding sequences of *Hydra vulgaris* Terb1, Terb2, and Majin using the predicted ORF sequences (Supplementary Fig. S7) from testis fraction. The predicted translation product from our full-length cDNA experiments showed similar domain organization with mouse TERB1, TERB2, and MAJIN protein sequence (Supplementary Fig.S8).

Taken together, our results are consistent with the notion that TERB1, TERB2 and MAJIN would fulfill meiosis-specific functions across metazoans.

## Discussion

### TERB1-TERB2 and MAJIN date back to the common ancestor metazoans

Active chromosome movements during meiotic prophase I is an evolutionarily conserved hallmark that ensures fidelity in the pairing of homologous chromosomes, synapsis and recombination. Despite the biological conservation of telomere anchoring to the NE, the sequence of meiotic telomere adaptor proteins is highly divergent between mouse and yeast. Our results casts new light on the evolution of the TERB1-TER2-MAJIN complex supporting the hypothesis of an origin that dates back to the common ancestor of metazoans.

In agreement with the commonly accepted view of basal metazoan phylogeny (Philippe et al., 2009), our analysis roughly recovers metazoan phylogeny separating non-bilaterians among Protostomia (Lophotrochozoa and Ecdysozoa) and Deuterostomia taxa despite the deepest nodes (BPP < 0.75) being were poorly resolved. In contrast, for the ecdysozoan clade only a few taxa belonging to Priapulida (*Priapulus caudatus*) and Arthropoda (*Zootermopsis nevadensis, Parasteatoda tepidariorum, Limulus Polyphemus, Centruroides sculpturatus*,) were identified as encoding homologous sequences. In addition, relatively long branches were computed indicating a fast evolutionary rate especially for Arthropoda. A possible explanation is that lineages leading to Nematoda and Hexapoda such as *C. elegans* and *D. melanogaster* (well-studied meiotic invertebrate model organisms) possess higher rates of divergence and gene loss in comparison to humans and cnidarians, which have greater gene similarity (Kortschak et al., 2003).

Overall, the existence of very distantly related sequences of TERB1, TERB2 and MAJIN in ancient non-bilateral clades such as Cnidaria, Placozoa and Porifera, seems likely that these genes are not an innovation of mammals. In this scenario TERB1-TERB2-MAJIN must have arisen only once in metazoan evolution and were then either secondarily lost or diverged beyond recognition in lineages leading to *C. elegans* and *D. melanogaster*. Our experimental detection and localization of orthologous transcripts in *Hydra vulgaris* testes strongly advocates a meiotic role for these genes in Cnidaria.

### TERB1, TERB2, and MAJIN share binding sites sequence domains across metazoans

Mouse TERB1, TERB2 and MAJIN proteins cooperate to establish the meiotic telomere complex by interaction with two other protein complexes: telomeric shelterin and the LINC complex (Shibuya et al., 2014; 2015; Long et al., 2017; Wang et al., 2017; Zhang et al., 2017; Dunce et al., 2018; Wang et al; 2019). TERB1 associates with telomeric DNA and binds both the TRF1 shelterin protein and TERB2-MAJIN. Our sequence analysis detected canonical ARM repeat and MYB-like domain organization of candidate TERB1 sequences in diverse phyla such as Chordata (Vertebrata), Arthropoda (Chelicerate), Mollusca (Gastropod and Bivalves), Annelida, Brachiopoda, *Hydra vulgaris* (Hydrozoa, Cnidaria) and *Amphideom queenslandica* (Porifera). Nevertheless, species that belong to Cnidaria (Anthozoa), Priapulida, Hemichordate, Cephalochordate and some Chordata (Vertebrata) possess TERB1 sequences that likely lost the MYB-like domain. The adaptable armadillo repeats structure enables diverse functions (Gul et al., 2017) and was suggested to interact with SUN1 (Shibuya et al., 2014;2015.). However, MYB domain or telobox consensus was specifically characterized to bind double telomeric DNA and associate with meiosis specific cohesion SA3 (Billaud et al., 1996; Shibuya et al., 2014; Daniel et al., 2014). The absence of a telobox domain in these phyla could imply sequence annotation errors. In addition, based on SCOP (Fox et al., 2013) classification the MYB-domain form part of the homeodomain-like super family’and is organized in three α-helices arranged in a helix-turn-helix motif that allows binding the DNA as a monomer, which is consistent with the secondary predictions of the multiple sequence alignment (Supplementary Information, Fig. S3).

Moreover, TERB1 localized at telomeres through the interaction with TRF1 (Shibuya et al., 2014). The TRF1-TERB1 association in mammals is through a peptide-docking sequence motif in human/mose TERB1 (aa 645-648) comparable to the strategy adopted by TIN2 (Chen et al., 2008, Long et al 2017). Despite it being demonstrated that mutations in this motif abolished or substantially impaired the binding of TRF1, leading to a decreased fertility in male mice (Long et al., 2017), our analysis failed to find this motif outside the clade of Vertebrata. This discrepancy could be attributed to a new adaption in vertebrate lineages to interact with TRF1. Further research should be done to characterize the mechanistic association and/or regulation in relation to the shelterin complex outside the clade of Vertebrata. Conversely, almost all TERB1 sequences analyzed bear the essential region for TERB2 binding (T2B). Recently, it was reported that the human TERB1 C-terminus (aa 590-649) interacts with the N-terminus of TERB2 (aa 2-116) through hydrophobic contacts and electrostatic interactions (Wang et al., 2019). In addition, the structure of T2B of TERB1 was resolved in two helices (H1 and H2) in which secondary structures alpha helix were nicely predicted for this region (Supplementary Information, Fig. S3).

Unexpectedly, our multiple sequence analysis identified a highly conserved motif in the N-terminal domain of TERB2 (aa 86-90 of human) in all taxa. This motif, resembles the [F/YxLxP] motif (Chen et al., 2008) required for shelterin proteins to bind the TRHF interface of TRF1 and TRF2, as previously discussed in mammals (Fig. 1B). These findings suggest that TERB2 could intrinsically be recruited to and/or interact with shelterin TRFH surfaces, further analysis needs to be done to confirm its functionality in the context of meiosis. In addition, the sequence alignment not only highlighted that the N-terminus of TERB2 was the most conserved region but also, detected a short stretch of similarity at the C-terminus which nicely correspond to the regions in which interact with N-terminus of MAJIN. Interestingly, a closer inspection of the identified MAJIN orthologues showed that the mainly conserved region is the N-terminus, while its C-terminal part is quiet variable in all taxa.

## Conclusion

Altogether, these results suggest that TERB1, TERB2 and MAJIN have an ancient origin in metazoan. Their detection in germ-line tissue of the ancient non-bilaterian *Hydra vulgaris* supports functional conservation over evolutionary time, implicating them as critical mediators of meiotic chromosome attachment.

## Material and Methods

### Data mining

The non-redundant protein database from the NCBI (https://www.ncbi.nlm.nih.gov/) was used to retrieve homologues of mice TERB1 (NP_851289), TERB2 (NP_083190), and MAJIN (NP_001159391) using PSI-BLAST (options: substitution matrix = BLOSUM45, word size = 3) (Altschul et al., 1997). Iterations were repeated for newly detected homologue sequences until convergence. Genomic sequences available at the NCBI (e.g. genome, est, tsa) and ensemble databases, were checked with the TBLASTN algorithm using the BLOSUM45 matrix. All retrieved sequences were used for reciprocal BLAST tests to ensure that they represented putative homologues of TERB1, TERB2, and MAJIN proteins and not false positives. Hydra vulgaris TERB1, TERB2, and MAJIN sequences obtained from this study (see below) were included in the analysis.

### Sequence alignment and phylogenetic tree construction

The retrieved sequences were aligned using PROMALS3D (Pei et al., 2008) which takes into account data from structural information for better understanding of sequence-structure-function relationships in distant homologues. Available structures from the Protein Data Bank (PDB) (Berman et al., 2003) for TERB1 (PDBI: 1×58_chainA and 6j07_chainB), TERB2 (6j07_chainA) and MAJIN (6j08_chainA) were uploaded in PROMALS3D. Annotations of the sequence alignment were designed using ESPript3 (Robert & Gouet 2014). Protein motifs and functional domains were examined through the Eukaryotic Linear Motif (ELM) (http://elm.eu.org/index.html) (Dinkel et al., 2016) and Superfamily web server (http://supfam.org/) (Gough et al., 2001) respectively.

For phylogenetic analyses, we selected a taxonomically balanced subset of homologous sequences which represent a wide range of animal phyla (Supplementary Information, Table S1). The sequences were aligned with MAFFT v7.309 with the accurate option L-INS-I. The resulting multiple alignments were trimmed with BMGE (“Block Mapping and Gathering with Entropy” software) v.1.12 (option -m BLOSUME45) (Criscuolo & Gribaldo, 2010). A Bayesian tree was computed with MrBayes v3.2.6 (Ronquist et al., 2012) with a mixed model of amino acid substitution including a gamma distribution (4 discrete categories) MrBayes was run with four chains for 1 million generations and trees were sampled every 100 generations. To construct the consensus tree, the first 2,000 trees were discarded as “burn in”. The final trees were drawn with iToL v4 (Letunic & Bork 2019).

### Maintenance of *Hydra vulgaris* strain AEP

Experiments were carried out using *Hydra vulgaris* strain AEP (Hemmrich et al., 2007). Animals were cultured according to standard procedures at 18°C (Lenhoff & Brown, 1970). Sexual differentiation was induced by starving the animals for several days after one week of intensive feeding.

### Isolation of RNA, reverse transcription, PCR, and cloning cDNA

Total RNA from *Hydra vulgaris* strain AEP, head, body column, testes and foot tissue were isolated using a peqGOLD TriFast kit (PeqLab) according to the manufacturer’s protocol. The RNA quality from different fractions was evaluated by non-denaturing agarose gel electrophoresis Subsequently, complementary DNA (cDNA) was synthesized from 1 μg of RNA, Oligo(dT)18 primer (Fermentas) and the M-MLV reverse transcriptase (Promega) according to the manufacturer’s instructions.

The cDNAs obtained were used as templates for the identification of the complete coding sequences and expression analyses in different tissues. To amplify full length coding sequence of *Hydra vulgaris* we designed specific primers using the predicted ORF sequences (Supplementary Information, Table S4). The cDNA amplification was performed with Phusion polymerase (Thermo-Fisher). PCR products were visualized on 1% agarose gel (peqLab) to verify size (Supplementary Information, Fig. S7), purified with a NucleoSpin Gel and PCR Clean-up Kit (Macherey-Nagel) and inserted into the pSC-A-amp/kan plasmid using a StrataClone PCR Cloning Kit (Agilent Technologies). The plasmids were purified using a NucleoSpin^®^ Plasmid kit (Macherey-Nagel) kit and verified by automated sequencing. To verify the obtained full-length or nearly full-length of cDNA, single read sequences from independent cloning were compared with the transcript reference sequence.

The expression studies were performed by RT-PCR using an intron spanning primer set (Supplementary Information, Table S4) designed the genomic annotations of Hydra vulgaris, TERB1 (Genbank ID: NW_004171015), TERB2 (Genbank ID: NW_0041710153) and MAJIN Genbank ID: NW_004173123). *Hydra* actin was used as a housekeeping gene to control for RNA amounts. The reactions were carried out with Phusion polymerase (Thermo-Fisher) for 32 cycles, so the amplification product was clearly visible. The products were visualized in a 1.2% agarose gel (PeqLab).

### Whole mount in situ hybridization

Antisense probes were generated by cloning a part of the putative *Hydra* homologues of TERB1, TERB2, and MAJIN (size between 400-700 bp) into the pSC-A-amp/Kan vector (StrataClone PCR Cloning Kit, Agilent) using specific primers (Supplementary Information, Table S4). After the validation of clone sequence, the vector was linearized for the synthesis of antisense RNA probes were performed by T7 and T3 RNA polymerase (Thermo-Fisher) with the incorporation of Dig-11-UTP (Roche Applied Science) at 37°C for 2 h. The RNA probes were purified using an RNeasy mini Kit (Qiagen) and checked by non-denaturing agarose gel electrophoresis. In situ hybridizations were performed according to the standard protocol described by Grens et al., 1996 with some modifications. Animals were relaxed in 2% urethane in hydra medium (HM) and fixed overnight with freshly made 4% formaldehyde in HM. The fixed animals were rehydrated and treated with proteinase K (0.05 mg/mL) for 20 min. To halt tissue digestion the animals were incubated with 2mg/mL glycine solution for 10 min. To minimize the background by preventing binding of the negatively charged RNA probe to positively charged amino groups on proteins samples were incubated in 0.1 M triethanolamine and acetic anhydride (0.25% and 0.5 % in 0.1 M triethanolamine) for 10 min each. The samples were re-fixed with 4% formaldehyde for 1 h at room temperature. Before hybridization the samples were washed with hybridization buffer (50% Formamide, 25% 20x SSC, 0.1% Tween 20, 0.15 mg/mL Heparin, 5mg/mL Torula RNA) at room temperature for 10 min and with pre-heated hybridization buffer at 57°C for 1 hr. Denatured digoxigenin (DIG) labeled probes were added to hybridization solution to a final concentration of 100 ng/mL, 20 h at 57°C. After several washes the samples were blocked in 1% blocking reagent in Maleic acid buffer MAB: 100 mM Maleic acid, 150 mM NaCl, pH 7.5) for 1 h at room temperature. For the detection of the DIG-labeled RNA probes an anti-DIG antibody coupled to alkaline phosphatase was preabsorbed 1:600 in (1X PBS, pH 7.5) on fixed animals overnight at 4°C and used directly in the same concentration at 4°C overnight. Unbound antibody was washed 15 times each for 20 min in 1X PBS, 0.1 % (v/v) Triton X-100. Detection of the signal was achieved through first equilibrating the animals with NTM (0.1 M NaCl, 0.1 M Tris-HCl, pH 9.5) for 10 min at room temperature and then incubated in 2% NBT/BCIP (Roche) in the dark. After reaching the optimal signal-to-background ratio, the reaction was stopped by washing the animals in ddH2O, followed by rehydration and mounting on microscop slides in 90% glycerol/1XPBS. The digital images were acquired using a in Leica EC3 digital camera incorporated on an Olympus SZ61 Zoom Stereo Microscope.

## Ethics statement

Animal care and experiments were conducted in accordance with the guidelines provided by the German Animal Welfare Act (German Ministry of Agriculture, Health and Economic Cooperation).

## Author contributions

I.d.C and R.B conceived and designed the study. I.d.C performed the identification of TERB1, TERB2 and MAJIN homologous sequences, multiple sequence alignment, modeling protein by structure homology and performed the expression analysis. C.B and I.d.C performed the phylogenetic analyses. I.d.C and R.B wrote the manuscript.

## Acknowledgements

We especially thank the DAAD (Germany) for granting I.d.C’s PhD fellowship. We also thank Thomas Bosch (Kiel) and his group for their support by providing *Hydra* cultures, Manfred Alsheimer (Würzburg) and José Sotelo-Silveira (Montevideo) for generous advice with the experimental design and for helpful discussions, Elisabeth Meyer-Natus (Würzburg) for excellent technical assistance, Isabell Köblitz (Würzburg) for advice with in situ hybridization techniques, Elena Bencurova and Brooke Morriswood (Würzburg) for critical reading of the manuscript. Supported by DFG grant Be1168/8-1 to R.B.

